# Marked *Neurospora crassa* strains for competition experiments and Bayesian methods for fitness estimates

**DOI:** 10.1101/736645

**Authors:** Ilkka Kronholm, Tereza Ormsby, Kevin J. McNaught, Eric U. Selker, Tarmo Ketola

## Abstract

The filamentous fungus *Neurospora crassa*, a model microbial eukaryote, has a life cycle with many features that make it suitable for studying experimental evolution. However, it has lacked a general tool for estimating relative fitness of different strains in competition experiments. To remedy this need, we constructed *N. crassa* strains that contain a modified *csr-1* locus and developed an assay for detecting the proportion of the marked strain using a post PCR high resolution melting assay. DNA extraction from spore samples can be performed on 96-well plates, followed by a PCR step, which allows many samples to be processed with ease. Furthermore, we suggest a Bayesian approach for estimating relative fitness from competition experiments that takes into account the uncertainty in measured strain proportions. We show that there is a fitness effect of the mating type locus, as mating type *mat a* has a higher competitive fitness than *mat A*. The *csr-1*\* marker also has a small fitness effect, but is still a suitable marker for competition experiments. As a proof of concept, we estimate the fitness effect of the *qde-2* mutation, a gene in the RNA interference pathway, and show that its competitive fitness is lower than what would be expected from its mycelial growth rate alone.

## Introduction

The filamentous fungus *Neurospora crassa* is a model eukaryote with a wealth of genetic resources (Roche *et al*., 2014; McCluskey *et al*., 2010; Colot *et al*., 2006), and many aspects of its cellular and molecular biology have been intensively studied (Roche *et al*., 2014). There is now a great deal of interest to study evolution of *Neurospora* and other filamentous fungi experimentally (Lee and Dighton, 2010; Graham *et al*., 2014; Romero-Olivares *et al*., 2015; Bastiaans *et al*., 2016; Fisher and Lang, 2016; Meunier *et al*., 2018). Despite having many beneficial characteristics for experimental evolution studies, *N. crassa* has lagged somewhat behind unicellular microbes in this area, as methodology for measuring competitive fitness has been missing.

Studying evolution of filamentous fungi is challenging because it is not clear how to define fitness in filamentous organisms (Pringle and Taylor, 2002). Many fungi have complicated life cycles, and individuals can be hard to define, complicating the choice of the appropriate fitness metric (Pringle and Taylor, 2002; Gilchrist *et al*., 2006). Moreover, individual fitness components, such as mycelial growth rate or conidial (asexual spore) production, are not necessarily strongly correlated with each other (Anderson *et al*., 2018). Yet, modeling results have shown that for saprotrophic fungi that colonize discrete resource patches, such as *N. crassa*, spore production is the critical fitness component (Gilchrist *et al*., 2006). While different experimental evolution protocols have been used for filamentous fungi, it has been shown that transferring spores to the next generation leads to the greatest response to selection (Schoustra *et al*., 2005). Accordingly, one should measure fitness in conditions that correspond to the propagation conditions. Therefore, spore production is often the measure of interest. However, just comparing spore production of two different genotypes does not necessarily predict which of the genotypes would prevail when the two are competing against one another. From studies with bacteria, we know that predicting the winner of two competing genotypes from their individual characteristics is difficult, and the best method is to measure competitive fitness directly (Lenski *et al*., 1998). Even maximum growth rate can be a poor predictor of fitness in bacteria (ConcepciÓn-Acevedo *et al*., 2015). Recently a study by Ram *et al*. (2019) showed that in some cases predicting competitive fitness from growth curve data is possible. However, this model would need to validated experimentally for *N. crassa*, and the model cannot take into account unexpected biological interactions between the strains. Therefore, competition experiments are still the best way to measure fitness.

To measure competitive fitness, one genotype needs to be tested against another genotype and the proportions of these genotypes in culture need to be followed. Following proportions requires that the genotypes are distinguishable. In controlled experiments, a morphological marker may have undesirable fitness consequences (Xiao *et al*., 2017), and often the genetic changes that happened between ancestor and derived genotypes are not fully known. Therefore, an engineered genetic marker is desirable. Some previous functional studies have used strains that express a fusion protein of histone H1 and green fluorescent protein to distinguish nuclei (Freitag *et al*., 2004) and different fluorescent labels could be used to distinguish between different strains. While using fluorescent labels is necessary for many functional studies, it requires imaging with a fluorescent microscope and counting of individual nuclei, which can be laborious in large evolution experiments. At the moment, a system to easily estimate competitive fitness of *N. crassa*, comparable to the *ara* marker in *Escherichia coli* where the *ara*^−^ and *ara*^+^ genotypes form red and white colonies when plated on tetrazolium arabinose agar plates (Lenski *et al*., 1998), does not exist.

To address the lack of suitably marked strains, we constructed genetically marked strains of *N. crassa*, and developed a PCR-based method to assess marker frequency in a sample of conidia. First, we introduced a DNA barcode by modifying the *cyclosporin resistant-1* (*csr-1*) gene, which encodes cyclosporin A binding protein. Mutations in *csr-1* gene cause resistance to the drug cyclosporin A, and this system allows replacing *csr-1* by homologous recombination (Bardiya and Shiu, 2007). Notably, this marker system allows the use of homologous recombination in *N. crassa* strains without disabled non-homologous end-joining DNA repair pathway (Ninomiya *et al*., 2004). Second, we used a high resolution melting (HRM) assay to distinguish between the amplification products of marked and wild type strains. HRM is a method in which a real-time PCR machine is used to monitor melting of PCR products. A fluorescent dye that binds double stranded DNA is included; when temperature increases, melting of the PCR products is monitored as the decrease in fluorescence caused by DNA strand separation. Sequence differences between different alleles cause their melting curves to differ, which can be used to distinguish them (Wittwer *et al*., 2003). By comparing unknown samples to known standards, the relative proportions of the different alleles can be determined. However, as with any other biochemical assay based on standard curves, there is some uncertainty associated with the standard curve and the samples. Therefore, we suggest a Bayesian statistical model to estimate fitness effects from competition experiments which incorporates all the uncertainty associated with our measurements. HRM has been previously used in several different applications, including genotyping (Wittwer *et al*., 2003), identification of different fungal species (Arancia *et al*., 2011), methylation analysis (Wojdacz and Dobrovic, 2007), and quantification of relative amounts of different bacterial strains in a sample to study competition (Ashrafi *et al*., 2017).

We show that our marked strain can be used in competition experiments, and that the HRM assay discriminates between the marked and the wild type strains. We further demonstrate that the marker itself has only a small fitness effect and illustrate the utility of our method by estimating competitive fitness effects for the different mating type idiomorphs, and for a mutant defective in the RNA interference pathway, *qde-2*, that is involved in environmental responses (Kronholm *et al*., 2016; Kronholm and Ketola, 2018).

## Materials and methods

### *Neurospora crassa* strains and culture methods

We used the nearly isogenic laboratory strains FGSC 2489 and 4200, obtained from the Fungal Genetics Stock Center (McCluskey *et al*., 2010), to generate a uniform genetic background. We backcrossed 4200 to 2489, always picking *mat a* offspring. Previously, we had performed five backcross generations (Kronholm *et al*., 2016), and now we performed further backcrosses until generation nine (BC_9_). From BC_9_ offspring we picked *mat A* and *mat a* genotypes to obtain BC_9_ 2489 *mat A* and BC_9_ 2489 *mat a*. Equivalent backcrossing was done for the *qde-2* mutant to obtain BC_9_ 2489 *mat A*; Δ*qde-2* and BC_9_ 2489 *mat a*; Δ*qde-2*. All strains used in this study, including the marked strains described below, are shown in Table 1. Hereafter we will refer to the BC_9_ 2489 background simply as 2489.

**Table 1:** Strains used in this study. Strains with genotype BC_9_ 2489 have the same genetic background generated by backcrossing 4200 nine times to 2489. FGSC = Fungal Genetics Stock Center, *wt* = wild type.

| FGSC ID | Genotype | Source |
| --- | --- | --- |
| 2489 | *wt mat A* | FGSC |
| 4200 | *wt mat a* | FGSC |
| B 26708 | BC<sub>9</sub> 2489 *mat A* | This study |
| B 26709 | BC<sub>9</sub> 2489 *mat a* | This study |
| B 26710 | BC<sub>9</sub> 2489 *mat A csr-1\** | This study |
| B 26711 | BC<sub>9</sub> 2489 *mat a csr-1\** | This study |
| B 26712 | BC<sub>9</sub> 2489 *mat A*; $\Delta qde-2$ | This study |
| B 26713 | BC<sub>9</sub> 2489 *mat a*; $\Delta qde-2$ | This study |
| B 26714 | BC<sub>9</sub> 2489 *mat A csr-1\**; $\Delta qde-2$ | This study |
| B 26715 | BC<sub>9</sub> 2489 *mat a csr-1\**; $\Delta qde-2$ | This study |

We used standard laboratory protocols to culture *N. crassa* (Davis and de Serres, 1970). Growth medium was Vogel’s medium N (Metzenberg, 2003) with 1.5% agar and 1.5% sucrose. Strains were grown in Lab companion ILP-02/12 (Jeio Tech, South Korea) growth chambers at 25 °C unless otherwise noted.

### Construction of marked strains

#### Making the *csr-1*\* construct

The overall strategy for making the construct used for transformation by PCR-stitching is illustrated in Figure 1. We made a linear construct that was homologous to *csr-1*, except that it contained a modified sequence (5’GAATTC<u>ATG</u>TAATAG-), which introduces an EcoRI restriction site (`GAATTC`), two early stop codons, `TAA` and `TAG`, and single base pair deletion causing a frameshift (Figure 1). The *csr-1* gene is on chromosome 1, between coordinates 7403946–7406381 on the reverse strand, while the modified sequence is located between coordinates 7405193–7405208 (*N. crassa* genome assembly NC12). We modified this part of the *csr-1* sequence because the `ATG` it contains is the initiation codon for the cytosolic isoform of the protein. A mitochondrial isoform is initiated from an alternative start upstream, and we wanted to abolish the function of *csr-1* completely.

**Figure 1:**
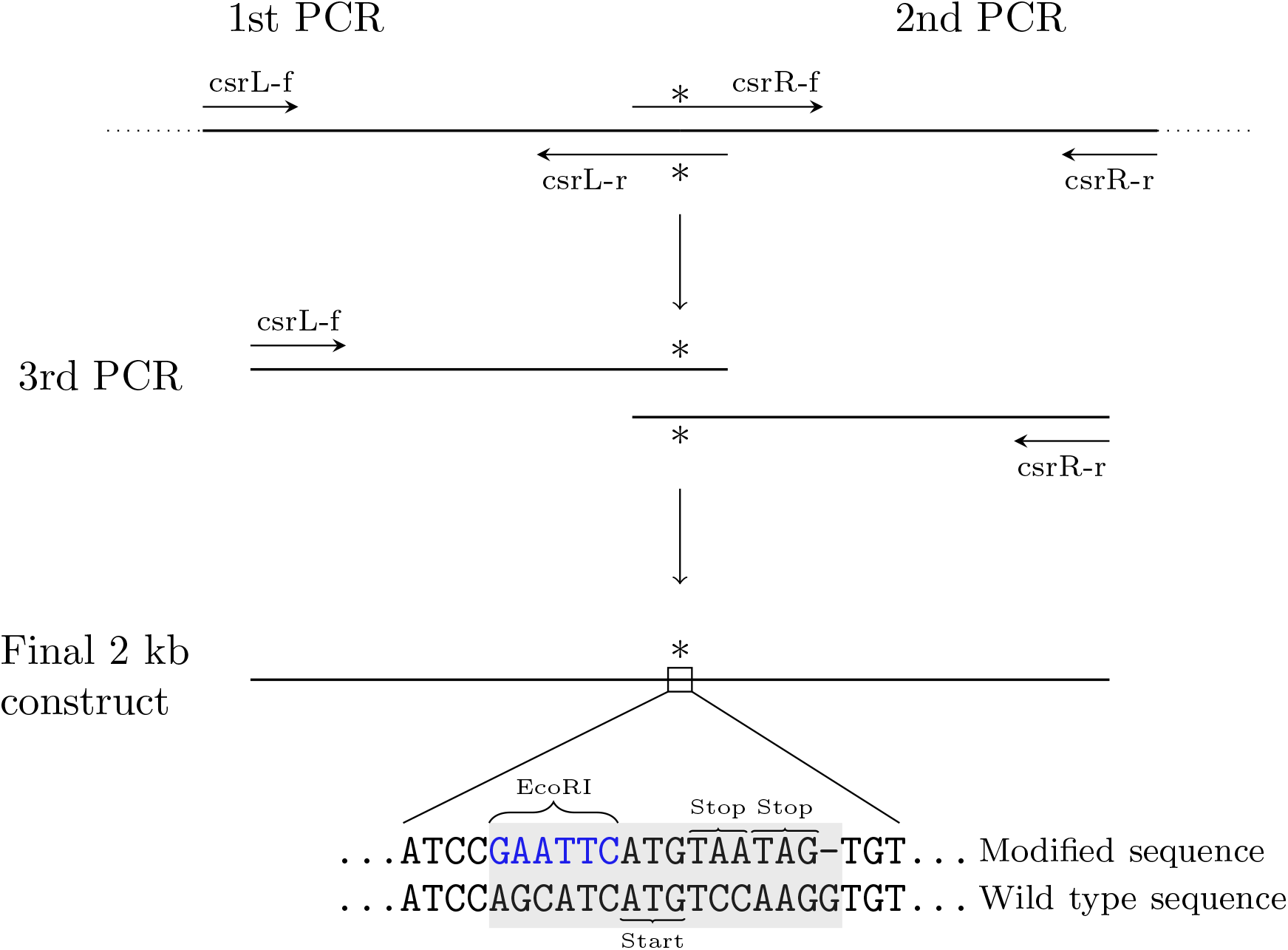
Overall strategy for generating the final construct by PCR-stitching. Primer names correspond to names in table S1. Sequence of reverse strand is shown, since *csr-1* is in the reverse strand.

To make the construct, we amplified two flanking 1 kb regions with primers such that one of the primers contained a tail with the new modified sequence and a region that was homologous to the other PCR-product (Figure 1). We used primers csrL-f and csrL-r to amplify the left flanking region and primers csrR-f and csrR-r to amplify the right flanking region (all primer sequences are given in Table S1). These products were amplified with the Phusion DNA polymerase (Thermo Scientific) in a 50 μl PCR reaction containing: 1 × HF-buffer, 0.2 mm each dNTP, 0.25 μM each primer, 100 ng of DNA, and 0.4 U of Phusion DNA polymerase. Reaction conditions for both reactions were: 1 min at 98 °C, then 30 cycles of 98 °C for 10 s, 60 °C for 30 s, 72 °C for 30 s, and a final extension at 72 °C for 5 min. PCR reactions were loaded on a 0.8% agarose gel and the 1 kb bands were extracted from the gel, and cleaned using GenCatch gel extraction kit (Epoch Life Science) according to the manufacturer’s instructions.

The final stitching PCR was performed using primers csrL-f and csrR-r with LA Taq polymerase (Takara) in a 50 μl reaction. The reaction contained: 1 × LA PCR buffer, 0.4 mm each dNTP, 2 μl of both csrL and csrR templates, 0.2 μM each primer, and 25 U of LA Taq. Reaction conditions were: 94 °C for 1 min, followed by 35 cycles of 98 °C 10 s, 68 °C 5 min, and final extension at 72 °C for 10 min. The PCR reaction was loaded on a 0.8% agarose gel, and the 2 kb band was extracted and purified as described above.

#### Transformation

We transformed the strain BC_9_ 2489 *mat A* by electroporation following Margolin *et al*. (1997). For electroporation, we mixed 200 ng of the construct and 40 μl of electrocompetent conidia in a chilled electroporation cuvette (2 mm gap), incubated on ice for 5 min, and electroporated with a Bio-Rad Gene Pulser using settings of: 600 Ω, 25 μF, and 1.5 kV. Immediately after electroporation, 950 μl of ice cold 1 m sorbitol was added to the cuvette and mixed.

Electroporated conidia were then transferred to a 50 ml conical tube with 9 ml of 32 °C liquid medium without sucrose, and incubated with shaking for 2 h at 32 °C. After the incubation, 10 ml of molten 2 × top-agar (standard growth medium with 2% agar, 2 m sorbitol, and 10 μg/ml cyclosporin A) was added to the culture and poured immediately onto selective medium (normal growth medium with 1.5% agar, and 5 μg/ml cyclosporin A). Cyclosporin A was dissolved in EtOH and added after autoclaving other medium components. Plates were incubated until colonies were visible, which were then picked and kept on slants.

#### Validation of strains

To confirm transformants, we screened candidates by PCR with csrL-f and csrR-r primers and EcoRI digestion to identify strains containing the modified sequence. We grew mycelia for each of the colonies in a 5 ml liquid culture for 2 days with shaking at 32 °C and harvested the mycelium, which was then lyophilized and pulverized. DNA was then extracted by a protocol adapted from Oakley *et al*. (1987). The original protocol was changed so that 1 ml of TCA/EtOH was used, and precipitation was done at −20 °C for 1 h. In addition, after incubation with RNase A solution (containing 0.15 mg/ml RNAse A), samples were extracted once with 200 μl chloroform and centrifuged for 5 min at 14 000 rpm. Supernatants were transferred to new tubes, and 900 μl of 8:1 isopropanol:7.5 m NH_4_OAc solution was added to each. Samples were mixed and centrifuged for 1 min, and supernatant was discarded. Pellets were washed once with 70% ethanol and air dried, and then resuspended in TE-buffer. 25 μl PCR reactions were performed using Phusion DNA polymerase as above with primers csrL-f and csrR-r. Cultures whose PCR product was digested with EcoRI were kept for further analysis. Sanger sequencing of positive transformants was done using standard protocols.

A positive transformant, BC_9_ 2489 *mat A csr-1*\*, was crossed to strain BC_9_ 2489 *mat a* to obtain BC_9_ 2489 *mat a csr-1*\*. Then BC_9_ 2489 *mat A*; Δ*qde-2* was crossed to BC_9_ 2489 *mat a csr-1*\* to obtain genotypes that were *csr-1*\*; Δ*qde-2* for both mating type idiomorphs. Progeny were screened by PCR using the high resolution melting assay for *csr-1* and PCR-protocol for *mat* locus and *qde-2* as in Kronholm *et al*. (2016).

### High resolution melting PCR

A high resolution melting (HRM) PCR assay of the *csr-1* gene was developed using primers (csr-hrm-f and csr-hrm-r, Table S1) that amplified a 100 bp PCR product containing the modified sequence. We amplified the product in 10 μl reactions containing 1×Precision melt supermix (Bio-Rad), 200 nm of each primer, and 2 μl of DNA. Sanger sequencing of the PCR product with primers csr-hrm-f and csr-hrm-r confirmed that the correct target was amplified. For HRM analysis, DNA was extracted by combining 40 μl of conidial suspension and 10 μl of extraction buffer (100 parts of 50 mm Tris pH 8 and two parts of 0.5 m EDTA pH 8.5) and incubating the mixture for 10 min at 98 °C in a PCR machine as in Kronholm *et al*. (2016). The same batch of buffer was used for all samples. PCR amplification was performed with a Bio-Rad CFX-96 Real-Time System PCR machine. The reaction conditions were: initial denaturation of 95 °C for 2 min, 40 cycles of 95 °C for 10 s, 60 °C for 30 s, and 72 °C for 30 s. Amplification was followed by a melting curve analysis: initial denaturation phase of 95 °C for 30 s and renaturation at 70 °C for 1 min, followed by a melt curve measurement from 70–90 °C by 0.1 °C intervals for 5 s. Fluorescence was monitored on the SYBR channel.

To construct standard curves for conidial mixtures that contained different known proportions of *csr-1*\* conidia, we grew the strains 2489 *mat A csr-1*\*, 2489 *mat a csr-1*\*, 2489 *mat A*, and 2489 *mat a* for 5 days on slants, and suspended the conidia in 1 ml of 0.01% Tween-80. We then measured conidial concentrations using a CASY TT cell counter (Roche) with a 45 μm capillary and a gating size range from 2.5 μm to 10 μm. We standardized concentrations to 10^8^ conidia/ml and combined different proportions of *csr-1*^+^ and *csr-1*\* conidia in 40 μl samples with proportions of *csr-1*\* conidia ranging from 1 to 0 by 0.1 decrements. We also tested a dilution series of conidia (from 10^8^ to 10^4^ conidia/ml) to asses the effect of starting concentration on HRM results. Competition experiment samples were run on 96-well PCR plates. To control for variation in PCR reaction conditions, we included two independently constructed standard curves on each plate, a no template control, and additional controls of pure *csr-1*^+^ and *csr-1*\* conidia.

### Competition experiments

#### Effect of mating type and *csr-1*\*

The first competition experiment had a fractional factorial design, in terms of *mat A, mat a, csr-1*^+^, and *csr-1*\*, where strains B26708–B26711 (Table 1) were combined in four strain combinations that had different *csr* alleles, as combinations where both strains have the same *csr* allele cannot be measured. We measured conidial concentrations as above, and standardized concentrations to 1 × 10^7^ conidia/ml for each strain to be able to pipet 10^5^ conidia easily. The experiment was started with 10^5^ conidia of both strains in a 75 × 25 mm test tube containing 1 ml of slanted agar medium. We repeated this experiment twice, first with five replicate populations for each competition, and then with 10 replicate populations for each competition, and we then combined these datasets. In total there were 60 populations. There were four time points, including zero, for each population, where the proportion of the *csr-1*\* was measured. Competitions were performed at 25 °C. Strains in the competition experiments were transferred every four or five days for three transfers. For a transfer, the conidia produced in a tube were suspended in 1 ml of 0.01% Tween-80 and 50 μl of conidial suspensions were transferred to new tubes. At each transfer, 40 μl of conidial suspension was taken for DNA extraction.

#### Fitness effect of the *qde-2* mutation

In the second competition experiment we used four strain combinations of strains B26708–B26715 (Table 1), so that competing strains always differed from each other in terms of mating type, csr, and *qde-2*. Five replicate populations for each different strain combination were used, giving 20 populations in total. Competitions were performed as described above, but at 35 °C for competitions with *qde-2* because in earlier studies we had observed that *qde-2* has an effect on growth at higher temperatures (Kronholm *et al*., 2016).

### Statistical analyses

All statistical analysis and data processing were done using the R environment version 3.5.2 (R Core Team, 2018). For Bayesian analyses we used the Stan language (Carpenter *et al*., 2017), that implements adaptive Hamiltonian Monte Carlo sampling. Stan was interfaced by the R package ‘rethinking’ (McElreath, 2015). For plotting we used the ‘ggplot2’ R package (Wickham, 2016).

#### Estimation of *csr-1*\* allele proportion

To process the melting curve data, we followed the approach used by Ashrafi *et al*. (2017) with some modifications. We first normalized the fluorescence data (RFU) between 0 and 1. The melting temperature for a given sample, i.e. the temperature of the inflection point of its melting curve, was found based on the maximum of the spline interpolation of the negative first derivative of the melting curve. To calculate the difference curve for the normalized RFU data, we subtracted RFU of the positive control of *csr-1*\* from the RFU of each sample. For further analysis using the normalized RFU differences, we used the temperature that gave the maximal differences between standard curve samples.

To estimate the proportion of conidia that contain the *csr-1*\* allele, we first built a standard curve and then estimated the proportion in unknown samples using this curve. For estimating the standard curve, we used the approach recommended by Ashrafi *et al*. (2017): the model was *y* = *a* + *Bx*, where *y* was the RFU difference or melting temperature and *x* was the proportion of *csr-1*\*, where *x* for unknown samples is estimated from

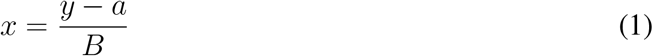

instead of fitting the proportion directly as a response. This approach allows fitting a normal distribution for the RFU difference or melting temperature, whereas proportion is constrained between 0 and 1. Thus, the Bayesian model for standard curve was:

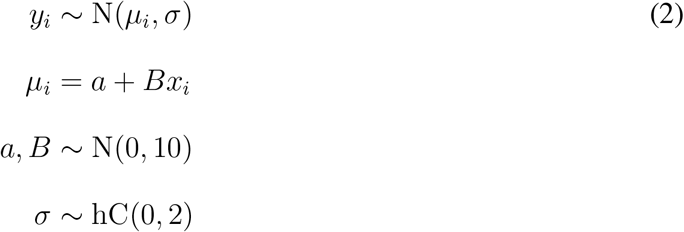

where *y_i_* was the *i*th observation of normalized RFU difference, *x_i_* is the *i*th *csr-1*\* allele proportion, *a* is the intercept, and *B* is the slope of the standard curve. We used weakly regularizing (McElreath, 2015) gaussian priors for *a* and *B*, and half-cauchy (hC) prior for the standard deviation *σ*. For MCMC estimation we used two chains, with warmup set to 1000 followed by 3000 iterations for sampling. Convergence of the model was examined by using trace plots and 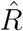 values, which were 1 for all estimated parameters. Proportion of unknown samples was estimated by using the posterior distributions for *a* and *B* and substituting them to equation 1. The values for proportion that were < 0 or > 1 were set to their limits. This way we obtained a posterior distribution for the *csr-1*\* allele proportion for each unknown sample.

#### Estimation of competitive fitness

Competitive fitness in haploid asexually reproducing organisms is just the ratio of growth rates of the competing types. Let *N* be population size, *r* growth rate, and *t* time, then population growth can be modeled as *N_t_* = (1 + *r*)^*t*^*N*_0_. If we have two competing types: *A* and *B*, then the ratio of these types grows as

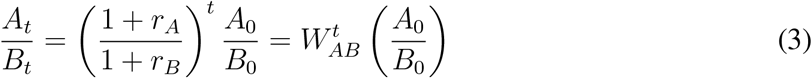

where *W_AB_* is the fitness of type *A* relative to *B*. We can replace the ratio of the absolute genotype numbers with the ratio of their proportions. Let *p* = *A*/(*A* + *B*) and *q* = *B*/(*A* + *B*), now *p*/*q* = *A*/*B* because the denominators cancel out. Taking a logarithm from equation 3 and substituting *q* = 1 – *p* yields

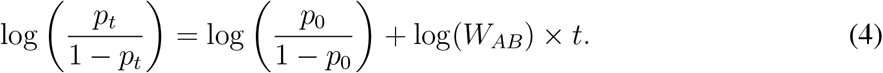

From this equation, we note that if we plot the log-ratio of the proportions of the two types against time, then log(*W*) is the slope of this line, and *t* the time of measurement. This is the standard way to estimate competitive fitness in asexuals (Hartl and Clark, 1997), and has been used extensively in the experimental evolution literature (Lenski *et al*., 1991). For *N. crassa*, this estimate works when strains are only allowed to reproduce asexually. We substitute the number of transfers for number of generations here; therefore, our fitness estimates include effects of mycelial growth rates and conidial production in as many cell divisions as it takes to go from spore to spore.

We included uncertainty in the *csr-1*\* allele proportion estimates in the model to estimate competitive fitness by using the observed posterior standard deviations for each *csr-1*\* allele proportion observation in the model (McElreath, 2015). To estimate relative fitnesses for the mating type and the *csr-1*\* allele, we fitted a model that accounted for population effects, effect of mating type, and the *csr-1*\* allele:

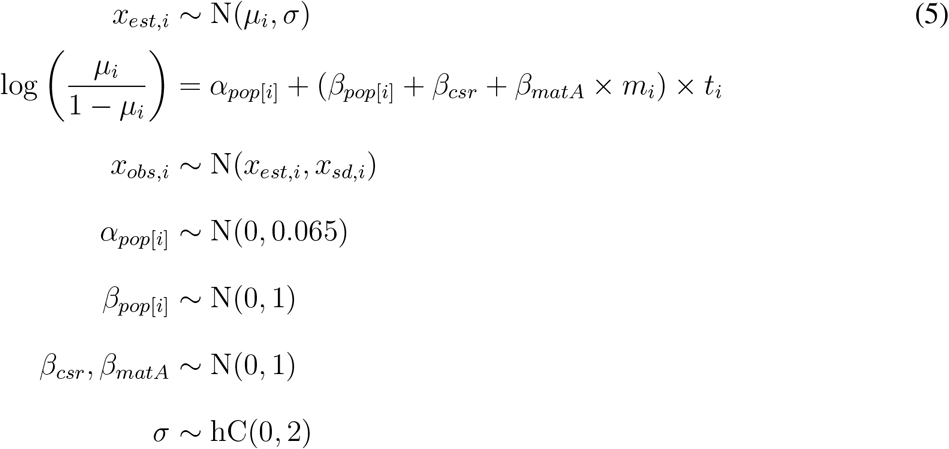

where *x_obs,i_* is the *i*th observed *csr-1*\* allele proportion, *x_sd,i_* is the observed error term for *i*th observation, *x_est,i_* is the estimated proportion for *i*th observation, *a*_*pop*[*i*]_ is the intercept for each replicate population (60 populations), *β*_*pop*[*i*]_ the slope effect of a replicate population (60 populations), *β_csr_* is the effect of the *csr-1*\* allele, *β_matA_* is the effect of mating type A, *m_i_* is an indicator whether in *i*th observation the *csr-1*\* allele containing strain is mating type A, *t_i_* is the transfer number for *i*th observation, and *σ* is the error standard deviation. Because these are competition experiments, where always two strains are competing, all the slope effects are relative effects, e.g. *β_matA_* is the fitness effect of *mat A* relative to *mat a*. Therefore, the indicator *m_i_* ∈ {−1, 0, 1}, so that *m_i_* = 1 when *csr-1*\* allele containing strain is *mat A, m_i_* = −1 when *csr-1*\* allele containing strain is *mat a*, and *m_i_* = 0 when mating types are identical. This way we can use all information in the data to estimate the effect of *mat A* from all competitions. We used weakly regularizing priors for the *β* slope effects, and an informative prior for the intercept, *α*. Since we started the competition with spores of both strains at a frequency of 0.5 or very close to this value, it makes sense to restrict intercept close to this value (0.5 is 0 on a logistic scale). The response of the model was fitted on the logistic scale, as at this scale relative fitness is the logarithm of the slope of this model, thus *W* = exp(*β*) for a given effect (Equation 4). MCMC estimation was done as above, but with 10000 iterations in total. Relative fitness of *mat A* and *csr-1*\* was calculated from posterior distributions of corresponding *β* effects.

When estimating the effect of the *qde-2* mutation, we first transformed the data such that the response indicated the frequency of the strain with the *qde-2* mutation and not necessarily the strain with the *csr-1*\* allele. The deterministic part of the model was:

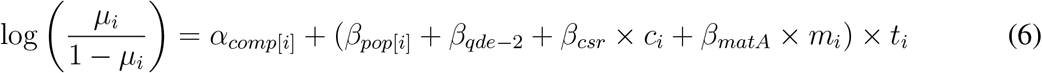

where *β_qde–2_* is the effect of the *qde-2* mutation, *β_csr_* is the effect of the *csr-1*\* allele, *c_i_* is an indicator variable whether in *i*th observation the *qde-2* strain has the *csr-1*\* allele, *c_i_* ∈ {−1, 1}, *β_matA_* is the effect of mating type A, and *m_i_* is an indicator variable whether in *i*th observation the *qde-2* strain is mating type A, *m_i_* ∈ {−1, 1}. Neither the *csr-1*\* nor the mating type effect are ever absent; they are just present in different configurations in the different competitions. *α_comp_* is the intercept effect for each competition (4 competitions). Other parameters, priors, and MCMC estimation were the same as above, but with 5000 iterations in total.

### Data availability

Strains generated in this study are available from the Fungal Genetics Stock Center (accession numbers B26708–B26715). The data and scripts implementing all statistical analyses are available from the University of Jyväskylä Digital Repository: https://doi.org/10.17011/jyx/dataset/65035

## Results

### Construction of marked strains

To introduce a marker to differentiate the strains in competition experiments, we modified the *csr-1* gene (ID: NCU00726) by homologous recombination. Rendering *csr-1* non-functional allows screening for positive transformants and distinguishing the strains by their *csr-1* sequences. After transformation we screened colonies for positive transformants; some of the colonies were heterokaryotic, but we found a homokaryotic transformant as well. We designated the new modified allele as *csr-1*\*. We subsequently validated the strains by Sanger sequencing, and observed the expected new *csr-1*\* and wild type *csr-1*^+^ sequences in a positive transformant and the wild type, respectively. We crossed the *csr-1*\* marker to different genotypes to have strains with both mating types and in the *qde-2* mutant background (Table 1).

### HRM assay optimization

To estimate the proportions of marked and unmarked conidia, we developed an HRM assay for the *csr-1* gene. We made mixtures with different proportions of *csr-1*^+^ and *csr-1*\* conidia. We could distinguish samples containing different proportions of the two *csr* alleles based on their melting curves (Figure 2A). The calculated melting temperatures were 81 and 82.1 °C for the *csr-1*\* and wild type alleles, respectively. However, for the 50% mixture, the melting temperature in our assays was 81.9 °C, which is not the midpoint between these two temperatures (Figure 2B). The alleles investigated here have multiple changes, so formation of heteroduplex DNA likely has a large effect on melting curve shape. Attempts to make standard curves with the melting temperature also yielded unsatisfactory results. Therefore, we used the difference in the normalized RFU instead (Figure 2C), picking the temperature where the difference between proportions of 0 and 1 were maximized to have the highest dynamic range. Using the normalized difference, we obtained good separation of the different proportions and a linear standard curve (Figure 2D).

**Figure 2:**
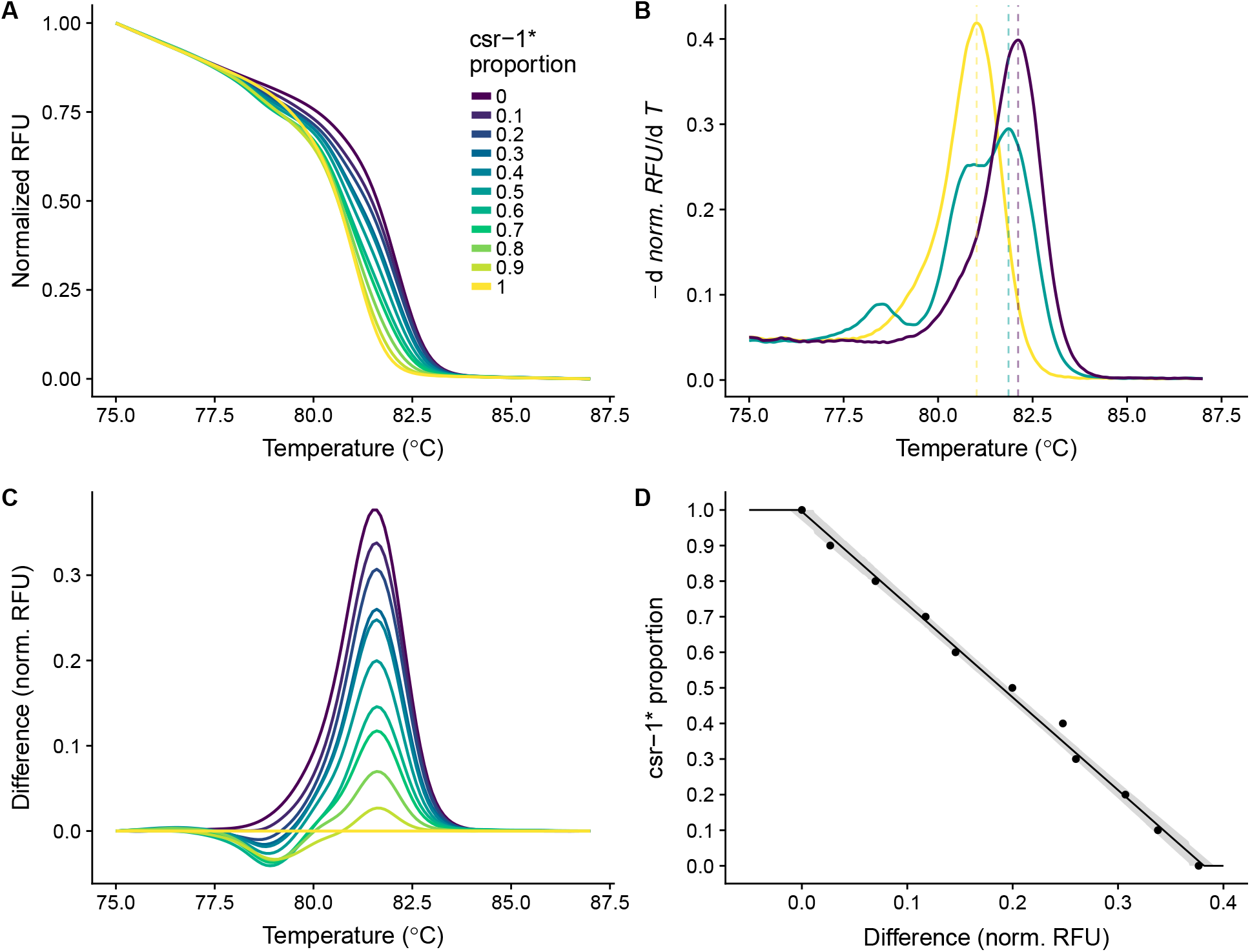
A) Melting curves of samples with different mixtures of *csr-1*^+^ and *csr-1*\* conidia. Normalized fluorescence against temperature. B) Melting temperature, i.e. the inflection point of the melting curve, can be found at the maximum of the negative first derivative of normalized RFU. Negative derivative against temperature for samples with *csr-1*\* proportions of 1, 0.5, and 0. C) Difference curves for the melting curves in panel A, relative to *csr-1*\*. D) Standard curve for *csr-1*\* allele proportion in the sample and normalized RFU difference.

To assess the efficiency of the PCR reaction, we tested the effect of the number of conidia in DNA extractions on the real-time PCR reaction. We observed that C_*q*_ value decreased with increasing numbers of conidia in the DNA extraction (Figure S1). The slope of this relationship was −1.9 when using log_10_ transformed number of conidia, and no significant differences were observed for the two alleles; the alleles amplified with similar efficiency. There are likely some PCR inhibitors in the conidial DNA extraction as the slope is > −3.3. However, inhibitors are unlikely to be a problem, since we are not interested in absolute quantification but rather relative proportions in one sample. Furthermore, the C_*q*_ values of competition experiment (see below) samples were similar; for the first competition experiment mean C_*q*_ = 26.0 and *σ* = 0.63, and for the second competition experiment mean C_*q*_ = 25.8 and *σ* = 2.35. The elevated standard deviation is due to a few samples having larger C_*q*_ values.

To summarize, the HRM assay allowed us to distinguish between *csr-1*^+^ and *csr-1*\* alleles and to estimate the proportion of these alleles in unknown samples using known proportions as a standard curve (Figure 2D).

### Competition experiments

Having a marker system that distinguishes strains from one another enabled us to perform competition experiments to estimate relative fitness of different genotypes. We inoculated two strains in one culture, and transferred conidia for three transfers and followed the strain frequencies using the HRM assay. This assay allowed us to estimate the relative fitness effects of different genotypes.

#### Effect of mating type and *csr-1*\* allele

In the first competition experiment, we tested the effect of the *csr-1*\* allele and the mating type locus. For the marker system to be useful, the marker itself should not have a large effect on fitness. We were also interested in the fitness effect of the mating type. When *N. crassa* mycelium grows, some hyphae fuse to form an interconnected network, which allows nutrient exchange within the mycelium. *N. crassa* strains that share the same mating type and heterocompatibility (*het*) alleles, such as the otherwise genetically identical strains used here, can fuse. Fusion is prevented for mycelia with different mating types (Metzenberg and Glass, 1990). In our experiments, we observed that when strains shared the same mating type, and thus fused with each other, the frequency changes of the *csr-1*\* allele were much smaller than with opposite mating types (Figure 3A). However, when mating types were different, and thus hyphal fusion was not possible, competitive exclusion seemed to happen quickly (Figure 3A). The *csr-1*\* allele had small frequency changes that seemed to go in both directions when mating types were the same (Figure 3A), indicating that the allele has no large fitness effects within the mycelium. For the 2489 *mat a csr-1*\* vs. 2489 *mat A* competition, the marked strain won in five replicates while the unmarked strain won in five replicates, and five replicates remained polymorphic (Figure 3A). In the 2489 *mat A csr-1\** vs. 2489 *mat a* competition, the 2489 *mat a* always won. Overall the estimate for the *mat A* fitness effect was: *W_matA_* = 0.58 (0.39–0.81, 95% HPDI) (Figure 3B). In the 2489 *mat A csr-1*\* vs. 2489 *mat A* competition, there were some frequency changes, but these were mostly small. However, in the 2489 *mat a csr-1*\* vs. 2489 *mat a* competition in some populations there was an initial larger change in frequency after which the frequency changes were smaller (Figure 3A). The effect of *csr-1*\* allele on fitness was negative: *W*_*csr*–1*_ = 0.71 (0.54–0.90, 95% HPDI). These results show that the mating type *mat a* has an advantage over *mat A* and that the *csr-1*\* allele has a fitness cost. It is likely that there is lot of variation in the outcome of the 2489 *mat a csr-1*\* vs. 2489 *mat A* competition because the marker effect cancels some of the fitness advantage of *mat a*, and in small populations chance has a larger role in determining the winner in this competition.

**Figure 3:**
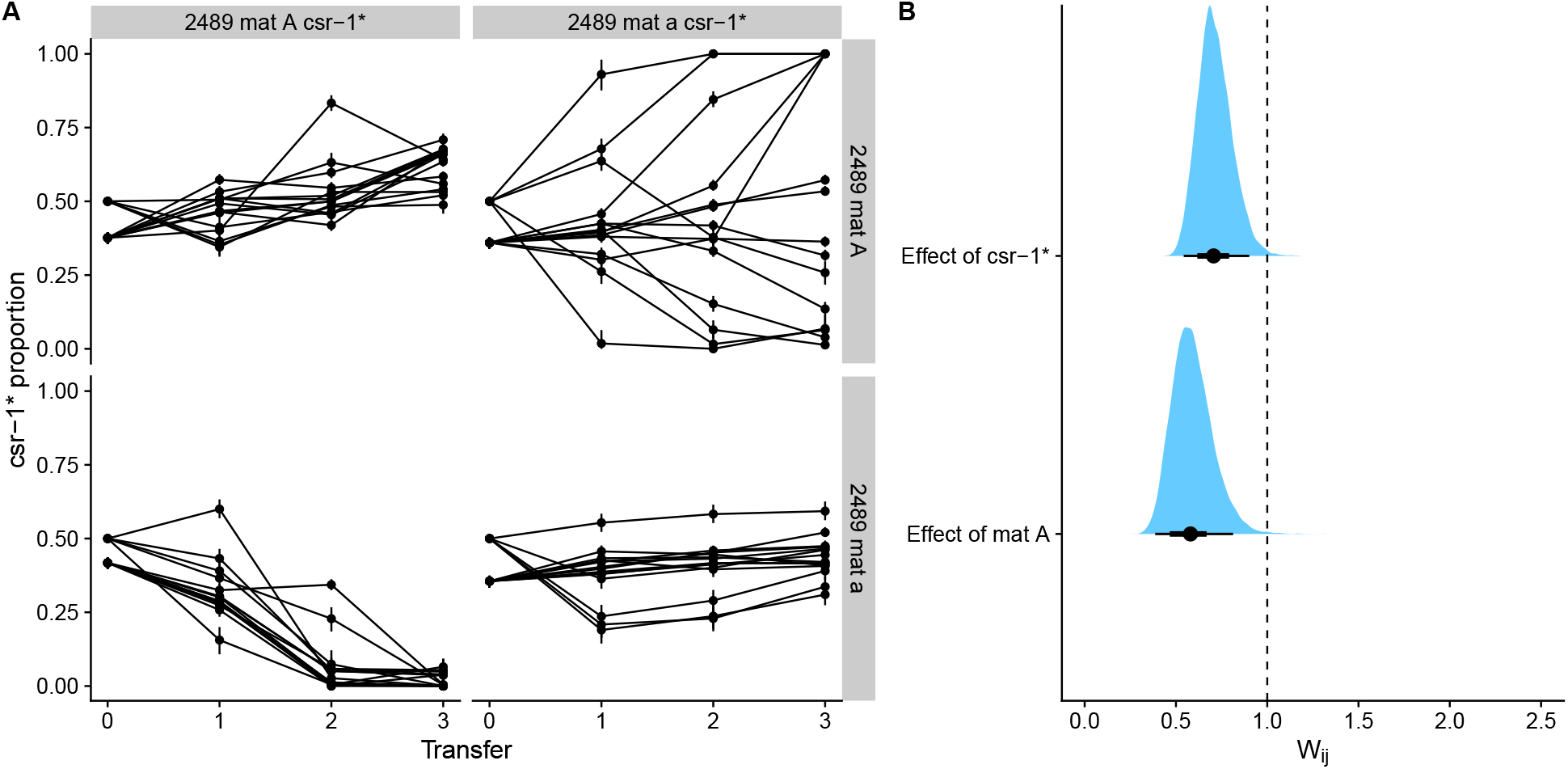
Results for the competition experiments testing the effect of the *csr-1*\* allele and mating type. For each treatment, *n* =15 for a total of 60 populations. A) Frequency trajectories of the *csr-1*\* allele. Some of the populations were started with slightly different initial proportions. B) Estimates of competitive fitness. Dots show point estimates for relative fitness, thick lines show the 66% and thin lines the 95% HPDI interval and the whole posterior distribution is filled above. The dashed vertical line shows relative fitness of one, i.e. no difference.

#### Fitness effect of *qde-2* mutation

Next we estimated the fitness effect of the *qde-2* mutation with competition experiments. QDE-2 (ID: NCU04730) is the *N. crassa* ARGONAUTE homolog and the corresponding mutant is deficient in small RNA processing (Maiti *et al*., 2007; Lee *et al*., 2010). We had previously examined the effect of *qde-2* on growth in different environments (Kronholm *et al*., 2016), and observed that it grows slower than the wild type particularly in temperatures of 35 °C and above. Therefore we expected that *qde-2* would have a lower relative fitness at 35 °C. Indeed, we observed that the strain with the *qde-2* mutation generally decreased in frequency (Figure 4A). In the 2489 *mat A* vs. 2489 *mat a csr-1*\*; Δ*qde-2* competition there was one population where the frequency of the *csr-1*\* allele initially decreased but then started to recover. Tthis phenomenon also happened, albeit to a lesser extent, for one population in the 2489 *mat A csr-1*\*; Δ*qde-2* vs. 2489 *mat a* competition (Figure 4A). The reason for this change of direction is unknown; one possibly is that a new beneficial mutation occurred in the *qde-2* background. Nevertheless, the *qde-2* strain clearly has a lower relative fitness compared to wild type (Figure 4B); the fitness estimate for *qde-2* was 0.30 (0.17–0.46, 95% HPDI). In this experiment, *mat A* had a suggestive effect: 0.68 (0.41–1.04, 95% HPDI), which is slightly higher than the effect in the previous experiment. Moreover, the fitness effect of *csr-1*\* overlapped with 1 in this experiment (Figure 4B), which is quite different than the estimate of the previous experiment. It may be that fitness effect of *csr-1*\* is so small that it is masked by the large effect of *qde-2*.

**Figure 4:**
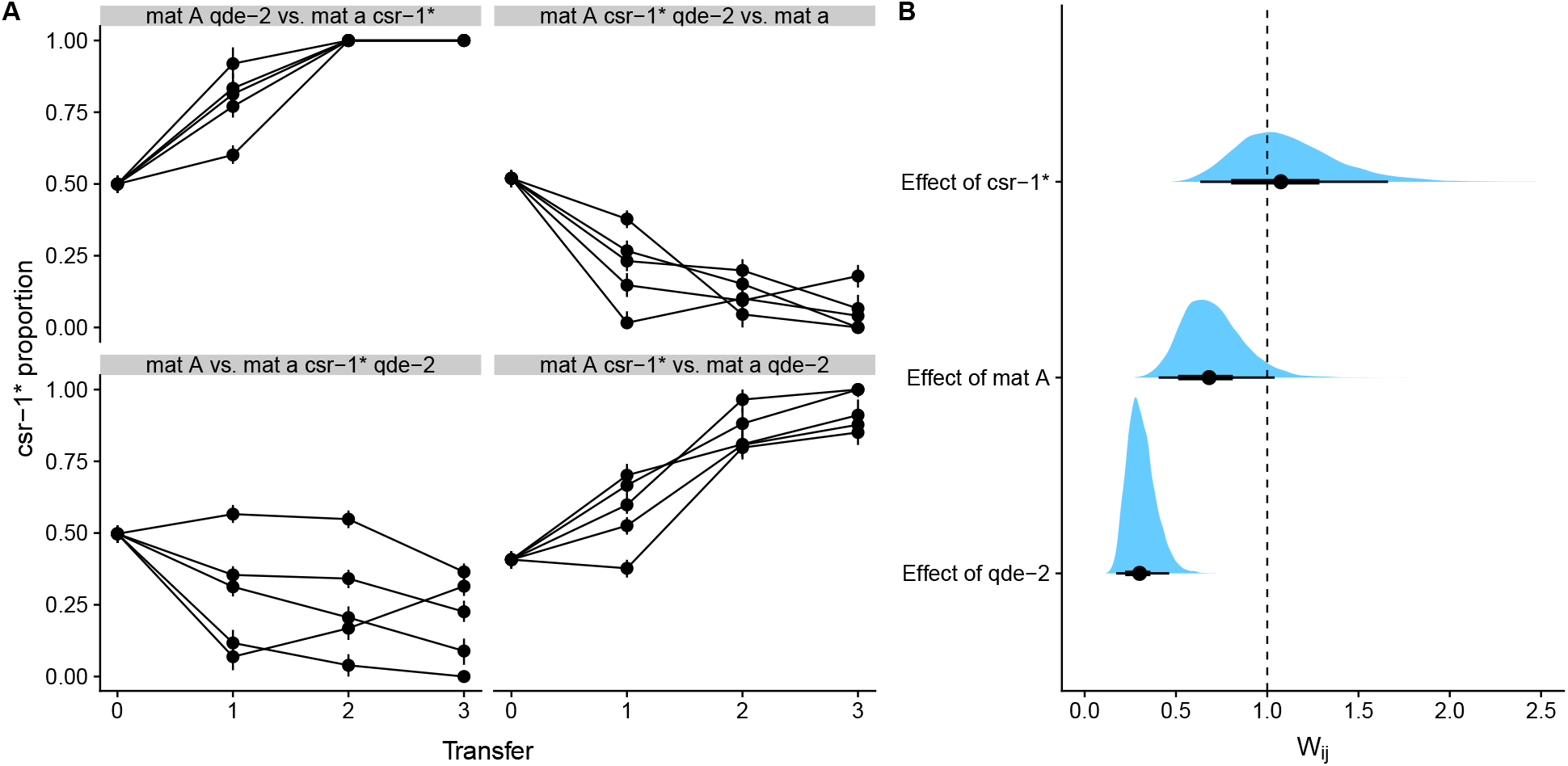
Results of competition experiments testing the effect of *qde-2* mutation. For each treatment, *n* = 5 for a total of 20 populations. A) Frequency trajectories of the *csr-1*\* allele. Note that raw data showing the frequency of *csr-1*\* is shown here, while fitness calculations were done with transformed data, where the frequency of the *qde-2* mutation was the response variable. B) Estimates of competitive fitness effects for *qde-2* mutation relative to wild type, mating type A relative to mating type a, and *csr-1*\* relative to *csr-1*^+^. Dots show point estimates for relative fitness, thick lines show the 66% and thin lines the 95% HPDI interval. The dashed vertical line shows relative fitness of one, i.e. no difference.

## Discussion

We showed that the *csr-1*\* allele can be used as a marker in competition experiments. The biological function of *csr-1* is not known, other than giving sensitivity to cyclosporin A (Bardiya and Shiu, 2007). The *csr-1*\* allele has a small fitness cost. However, both the effect of the marker and a genotype of interest can be estimated separately if the experimental design includes swapping the marker between the competing strains as done here. If swapping experiments are not possible, the known effect of the marker could be included in the model via priors. Considering that the marker has a small fitness effect, we recommend that experimental design include marker swapping experiments whenever possible. In this way the *csr-1*\* allele is a suitable marker for quantification of competitive fitness in *N. crassa*.

We also observed that when the two strains had the same mating type, the *csr-1*\* allele frequency changed only slowly or was maintained close to 0.5, but when strains of two different mating types were competing there seemed to be competitive exclusion of one of the strains. In *N. crassa*, genetically compatible strains undergo hyphal fusion only between strains that have the same mating type and compatible heterokaryon incompatibility alleles (Metzenberg and Glass, 1990; Zhao *et al*., 2015). Since our strains are nearly isogenic, they can undergo hyphal fusion and form a heterokaryotic mycelium with respect to the *csr-1* locus, maintaining both alleles. Fusion likely happened in competitions of strains with the same mating type. Nuclear ratios in *N. crassa* heterokaryons seem to be determined at the establishment phase of the heterokaryon, and the ratio of nuclei can be rather stable afterwards (Atwood and Mukai, 1955; Pittenger and Atwood, 1956). Frequency changes of different nuclei apparently require that the nuclei have different rates of mitosis, as diffusible components seem to be shared within the mycelium (Pittenger and Atwood, 1956). In the related species *N. tetrasperma*, which has a pseudohomothallic mating system, there is some evidence that the different nuclei are maintained by active processes (Samils *et al*., 2014). Furthermore, in the basidiomycete *Heterobasidion parviporum* ratios of different nuclei are affected by genetic and environmental effects (James *et al*., 2008).

In contrast, when strains with different mating types were competing, there seemed to be competitive exclusion, in which one strain often came to dominate the culture. When strains with different mating types fuse, cell death occurs in the fused cells, and these strains are thus unable to form heterokaryons (Metzenberg and Glass, 1990), and competition must happen. Based on theoretical modeling of fungal fitness for a filamentous fungus life cycle in a system of many habitable patches, competitive exclusion between different strains is inevitable (Gilchrist *et al*., 2006). In our experiments, the strains were competing only for a single patch. In the laboratory, *N. crassa* seems to follow a bang-bang life history strategy (Gilchrist *et al*., 2006), where the mycelium first grows to cover nearby habitable area and then the fungus switches to spore production. Hence, our strains may be competing mainly for space in the culture tube. In some populations of the mating type and *csr-1*\* allele experiments, the mating type A seemed to win even if overall *mat a* had a higher fitness. In these cases, it may be that *mat A* gained some initial advantage due to chance, and thus gained an advantage by simply having more spores in the next transfer. During *N. crassa* spore germination, spore fusion increases growth (Richard *et al*., 2012). Therefore, a strain with more spores, and thus more potential fusion partners, may have an initial advantage even if its steady state growth rate of mature mycelium is slightly slower.

Surprisingly, we observed a fitness effect for the mating type locus with *mat a* having a higher relative fitness than *mat A*. In a facultative sexual species such as *N. crassa* both mating types have to be maintained in a population for sexual reproduction to occur; so it seems unexpected that one mating type has a higher asexual fitness than the other. However, it has been reported previously that in *N. crassa mat a* has a higher growth rate than *mat A* (Ryan *et al*., 1943), and that germination of *mat A* conidia is slightly slower (Wang *et al*., 2019). This difference can be understood in the light of gene expression differences, as many genes unrelated to sexual reproduction are expressed differently in the vegetative mycelium of the different mating types, despite that these lines are nearly isogenic except for the mating type locus (Wang *et al*., 2012). Thus, the mating type genes can influence growth and other phenotypes. Different mating types have been reported to have different growth rates also in other species: including *N. tetrasperma* (Samils *et al*., 2014), *Pleurotus ostreatus* (Larraya *et al*., 2001), and *Fusarium culmorum* (Irzykowska *et al*., 2013). Considering that mating types can have differential fitness, how are they maintained in natural populations? One possibility is that the observed fitness advantage could be environmentally dependent, or alternatively, that sexual reproduction occurs often enough that mating type frequencies approach the evolutionary stable ratio of 1:1 despite differences in asexual fitness.

The *qde-2* mutation was known to grow slower than the wild type at 35 °C (Kronholm *et al*., 2016), so the observation that it also has a lower competitive fitness is expected. The results showed that we can measure fitness effects of mutations with our marked strain system. The magnitude of the fitness effect is perhaps larger than expected: the growth rate of *qde-2* is 79% that of wild type at 35 °C (Kronholm *et al*., 2016), while the relative fitness of *qde-2* was 30% that of wild type. Thus, the relationship between mycelial growth rate and competitive fitness is not a simple one to one relationship. Similarly, we have not observed large effects on growth rate for the *mat* locus, even if *mat A* has lower fitness. Moreover, the estimates seem large when compared to fitness effects of gene deletions in yeast (Bell, 2010). One possible explanation is that these estimates are not directly comparable as there is a large number of mitotic cell divisions between transfers, from spore to spore, and one transfer in a filamentous fungus is not comparable to a cell division generation in unicellular yeasts or bacteria.

The *csr-1*\* allele is a very versatile marker. In this study, we used HRM PCR as a method to detect the proportion of the *csr-1*\* allele, but there are other methods to estimate *csr-1*\* allele proportion in a sample of spores. The simplest, although more laborious, method is to do replica plating of spores on plates with and without cyclosporin A. Other PCR-based methods besides HRM that can detect the sequence difference between the *csr-1* alleles could also be used to estimate the proportion of the *csr-1*\*, such as digital droplet PCR with different probes (Hindson *et al*., 2011), pyrosequencing (Harrington *et al*., 2013), or any next generation sequencing technology, where many samples can be multiplexed with different barcodes and sequenced together (Smith *et al*., 2010). The advantage of our method is that it only requires to set up conventional PCR reactions taking little bench time, and the whole process can be done in 96-well plates enabling moderate throughput.

Another advantage of our method is the Bayesian model employed. Standard curves are commonly used in various biochemical assays, and while Bayesian approaches have been developed for various assays (Gelman *et al*., 2004; Feng *et al*., 2010) they are rarely used in practice. Our model allows any uncertainty arising from the standard curve, the samples, or the initial inoculation via measuring proportion at transfer 0, to be propagated into the fitness estimates. Our modeling approach could also be used for other marker systems employing standard curves.

In conclusion, the marked strains reported here can be used to measure fitness effects of individual mutations or even for the fitness of strains derived by experimental evolution. They provide a versatile tool and advance the use of *N. crassa* as system studying experimental evolution and ecology (Lee and Dighton, 2010; Fisher and Lang, 2016).

## Acknowledgements

This study was supported by the Academy of Finland grant no. 274769 to IK and no. 278751 to TK. KJM was supported by National Institutes of Health Training Grant: T32 HD007348. EUS was supported by NIH grants GM093061 and GM127142. We thank Matthieu Bruneaux, Roghaieh Ashrafi, and Neda Moghadam for comments on the manuscript. We also thank two anonymous reviewers for their constructive comments that improved the manuscript.

## Supplementary Information

### Supplementary Figures

**Figure S1:**
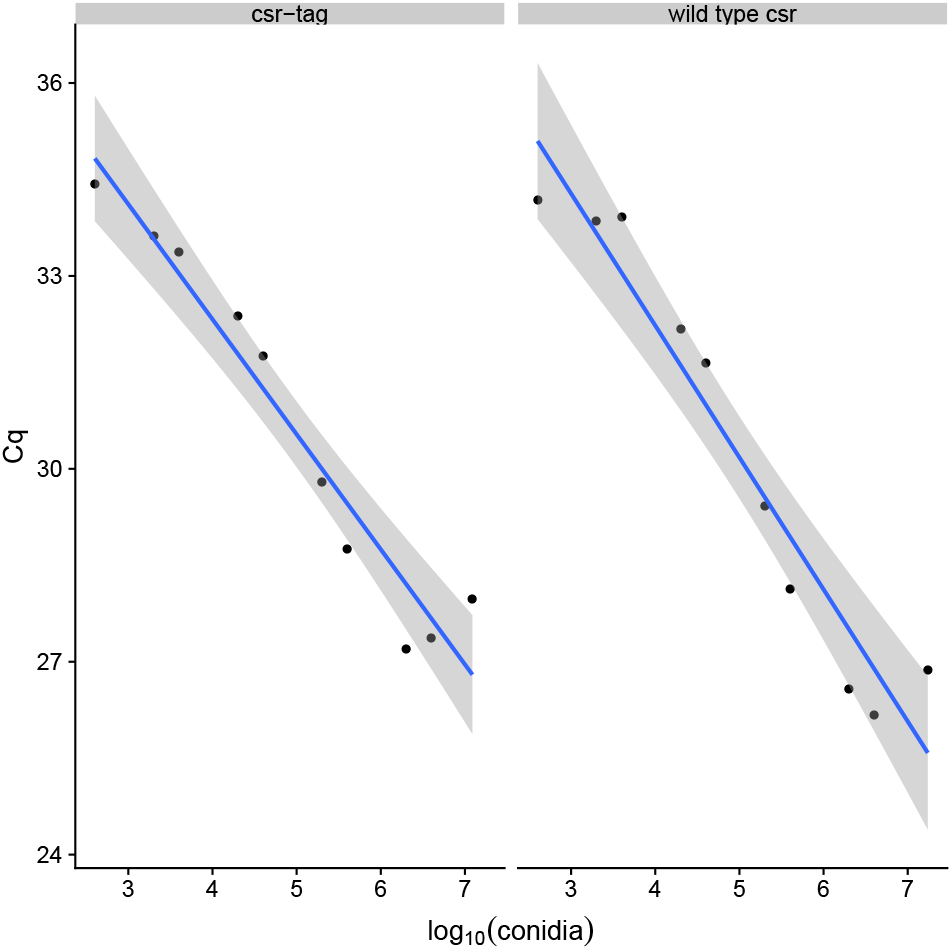
Correlation between number of conidia used in DNA extraction (log-scale) and number of PCR cycles required to detect the PCR-product. Differences between the alleles are not significant.

### Supplementary Tables

**Table S1:**
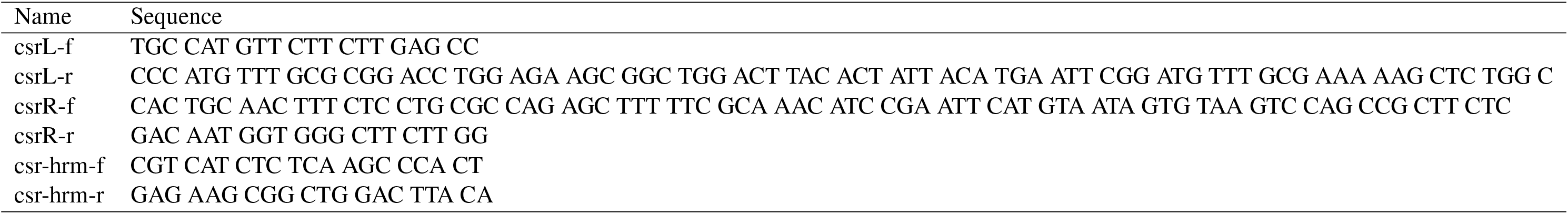
Primers used in the study.

## Notes

#### Summary of Updates

We have revised the manuscript according to reviewer suggestions. In particular we have collected new data to increase the sample size for the experiment where we estimate the fitness effects of the mat locus and the csr-1* marker. See results subsection: "Effect of mating type and csr-1* allele" and Figure 3. Our estimates are now more precise, and there is now also a significant effect of the marker. Manuscript has been revised accordingly. However, our method is still useful as one can perform marker swapping experiments to estimate marker effects and the effect of genotype of interest. Alternatively the effect of the marker can be included in the model via priors. Various other clarifications have been made, and typos have been corrected.

